# Century-old ethanol-preserved Vega Collection reveal the unexpected phylogeography of Slender bitterling *Tanakia lanceolata* in Japan

**DOI:** 10.1101/2025.11.02.686072

**Authors:** Keisuke Tsuchiyama, Seigo Kawase, Yuko Takigawa, Tomohiko Fujita, Kazumi Hosoya, Jyun-ichi Kitamura, Tappei Mishina

## Abstract

Recent genetic advancements offer new opportunities for studying natural history museum collections. The genetic analysis of century-old aquatic animal specimens, mostly preserved in ethanol, can provide valuable insights into the changes in genetic diversity caused by anthropogenic impacts. However, knowledge of the characteristics of degraded DNA from such specimens remains limited. In this study, we evaluated the DNA quality of bitterling fish, *Tanakia lanceolata* (Temminck and Schlegel), collected during the Vega Expedition in Japan in 1879 and preserved in ethanol. We then performed genomic analysis to test its hypothesized translocation-involved population history. The historical DNA was degraded, peaking around 50 bp, but a minor fraction exceeded 300 bp. Mitochondrial genetic analysis revealed high genetic similarity between the eastern and western sides of the Central Highland, typically impeding the dispersal of primary freshwater fish. These findings suggest an unexpected dispersal ability in *T. lanceolata* or undocumented translocations before the large-scale domestic translocations recorded since the 1910s. The customized workflow for historical ethanol-preserved specimens, based on historical DNA quality in this study, provides a foundation for studying historical archives.

## Introduction

Natural history museum collections are invaluable archives of global biodiversity preserved with detailed records, including collection dates and locations (Dawkins 1877). While these specimens were historically deposited for morphological studies, recent advancements have offered opportunities for genetic research (Sproul and Maddison 2017; Raxworthy and Smith 2021). These collections are particularly vital for conservation genomics, as they help develop evidence-based conservation strategies, an emergent approach called “conservation museomics” (Benham and Bowie 2022; Jensen et al. 2022; Blair 2024). Accessing DNA from past populations has helped resolve phylogenetic relationships (Ruane and Austin 2017; Orr et al. 2021), infer historical biogeographic structure (Martin et al. 2014), and clarify taxonomic confusion with type specimens (Kehlmaier et al. 2019; Cong et al. 2021). Additionally, comparing between historical and modern populations offers insight into anthropogenic impacts, such as extinction, population decline, and translocation (Valk et al. 2019; Roycroft et al. 2021; Andrews et al. 2025).

Specimens collected over a century ago are particularly valuable, as they are likely less affected by widespread anthropogenic impacts on biodiversity. Nevertheless, such specimens could provide crucial snapshots of the long-term human impacts. Modern aquatic animal collections are fixed in ethanol (typically 70%–80%) or formalin. Since formalin fixation became well established in the early 1900s (Ferdinand 1893; Fox et al. 1985), ethanol-fixation was preferred before that period. The historical DNA (hDNA) of these ethanol-preserved specimens is expected to be degraded during long-term storage at room temperature, especially when specimens have experienced low ethanol concentration due to evaporation during long-term preservation. Despite the challenges posed by degraded hDNA, several studies have successfully applied phylogenetic analysis to century-old museum collections preserved in ethanol, including snakes (Ruane and Austin 2017; Zacho et al. 2021), lizards (McGuire et al. 2018), and fishes (Silva et al. 2019). However, optimal conditions for analytical success in such old specimens remain unclear (Raxworthy and Smith 2021), as few studies have systematically assessed hDNA quality relative to specimens conditions. This can result in unexpected analytical difficulties and limit access to valuable historical collections. Accordingly, establishing an experimental framework that considers both hDNA quality and specimen condition is essential for the effective use of these valuable specimens.

Among the specimens collected over a century ago, a valuable example is the Vega Collection in the Swedish Museum of Natural History, Stockholm. The Vega was the first ship to navigate the Arctic Ocean route across the Eurasian continent, the shortest path between Asia and Northern Europe, from 1878 to 1879. After completing the Arctic voyage, the ship made port calls in Japan (Yokohama, Kobe and Nagasaki). Nils Adolf Erik Nordenskiöld (1832–1901), the leader of the Vega Expedition and a geologist, collected various animals, fossils, and minerals during the journey. Freshwater fishes were primarily collected from the Kanto region and Lake Biwa in the Kinki region, Japan (Fig. 1A) (Fujita 2019). Nordenskiöld recorded in his journal that he preserved the fishes in alcohol (Nordenskiöld 1880). Although the collection has yet to be investigated, a preliminary review of the freshwater fish collection from Lake Biwa consists of 15 species and 73 individuals in 25 lots from at least six sites (Takigawa et al. 2020). These rich specimens provide an opportunity to reconstruct historical genetic profiles and investigate changes in fish fauna over more than a century.

**Figure 1.**
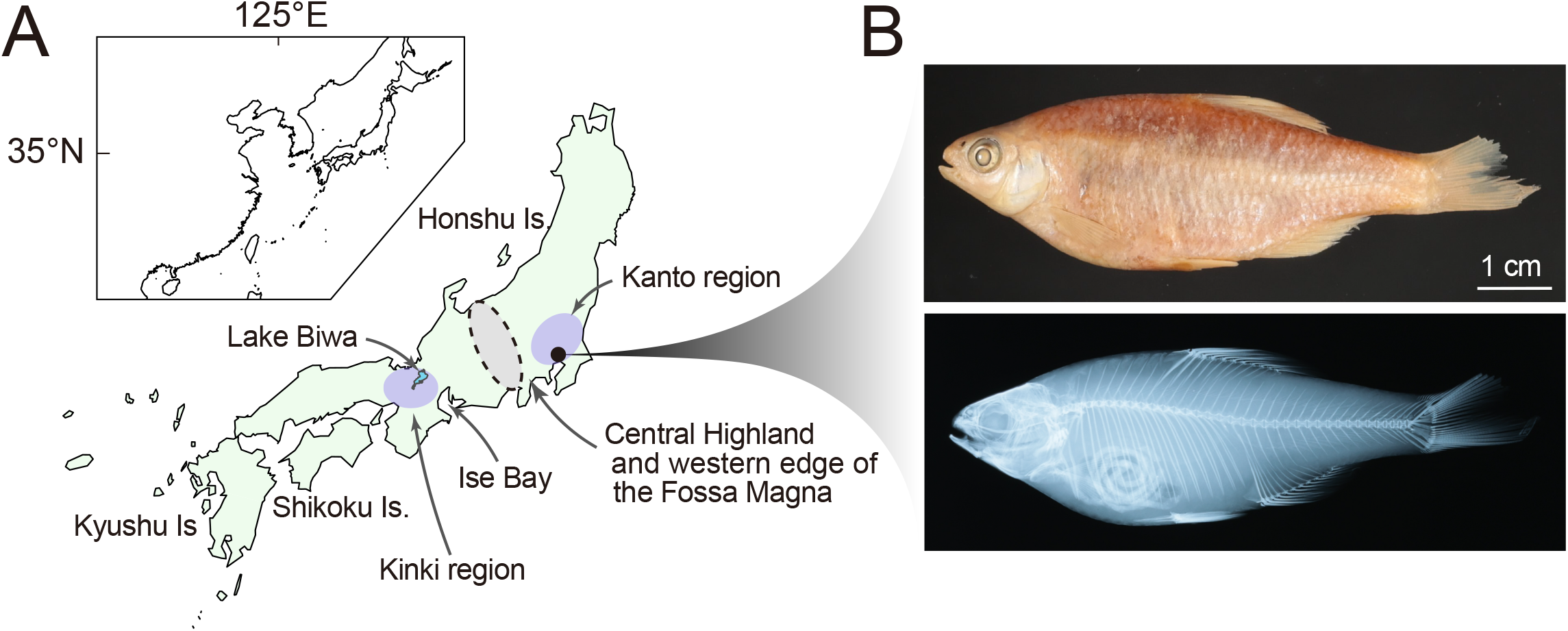
Discovery of *T. lanceolata* in ethanol-preserved Vega Collection from the Kanto region in 1879. (A) Map of the sampling site and related geological information. The black dot indicates the most likely registered collection site, “Yokogawa River near Tokyo”. A minor possibility remains that the collection originated from the Yokogawa River in Gunma Prefecture, also in the Kanto region along the recorded land excursion route to Mt. Asama, but far from Tokyo. (B) Lateral view (top) and soft X-ray image (bottom) of a representative *T. lanceolata* specimen reidentified from the Vega Collection.

A compelling model for this investigation is the Slender Bitterling, *Tanakia lanceolata* (Temminck and Schlegel), a primary freshwater fish widely distributed across Honshu, Shikoku, and Kyushu islands in Japan, as well as the Korean Peninsula and the northern part of China (Kitamura and Uchiyama 2020). Examination of the Vega Collection revealed bitterling specimens from the Kanto region with morphological traits consistent with *T. lanceolata* (Fig. 1B, Fig. S1A, B). These specimens offer an opportunity to investigate hypothesized extensive domestic translocations in the area. A recent comprehensive phylogeographic study using mitochondrial DNA revealed that some Kanto populations are genetically indistinguishable from those in the Kinki region (Tominaga et al. 2020), deviating from the typical east–west divergence observed in many freshwater fishes due to the geological barrier by the Central Highland in the central Honshu (Fig. 1A). Thus, a previous study suspected that the irregular distribution of mitochondrial haplotypes in this species across the Kinki and Kanto regions results from the translocation from the Kinki to Kanto (Tominaga et al. 2020), causing the mitochondrial introgression into local populations or questioning the existence of truly indigenous lineages.

This study evaluated the preservation quality of specimens collected during the Vega Expedition, assessed the integrity of their 140-year-old hDNA, and performed a genomic analysis to test the suspicious translocation-involved population history of *T. lanceolata* in the Kanto region. Our assessment revealed that the hDNA was degraded, but retained a notable peak around 50-bp, with fragments extending beyond 300-bp, allowing efficient use for genomic analyses. The mitochondrial genetic characteristics of *T. lanceolata* from the Vega Collection were very similar to those of modern specimens.

These results suggest an unexpected dispersal ability in *T. lanceolata*, which may complicate interpretations of its translocation history or suggest previously unrecognized historical translocations of this fish before 1879. The customized workflow established here, tailored to the hDNA quality of historical ethanol-preserved collections, provides a foundation for future studies using museum specimens.

## Materials and Methods

### Preservation condition and morphological identification of the Vega specimens

Studied material, collected during the Vega Expedition, is part of the wet collection at the Swedish Museum of Natural History (NRM). Since 1996, the wet collection has been kept in a dark storage room at a temperature of 15–18 °C throughout the year. Before the establishment of this storage room, the temperature was more variable and most certainly a few degrees above 20 °C during the summer. We reidentified 44 specimens from NRM 8168 (collected on Oct. 8, 1879, from the Yokogawa River near Tokyo, Japan) registered as *Acheilognathus melanogaster* Bleeker (Fig. S1C, D). The morphological identification was based on Hosoya (2013). Notably, 43 out of 44 specimens were identified as *T. lanceolata* (re-registered as number of NRM 68053), and morphological characteristics of 30 *T. lanceolata* and 1 *A. melanogaster* specimens were examined precisely (see “Result and Discussion”; Table S1). Measurement and counting methods followed Hubbs and Lagler (2004), and all measurements were taken to the nearest 0.1 mm using a digital caliper. The last two rays of the dorsal and anal fins were counted as one ray. Soft X-ray radiographs were taken to observe the bone condition and count the number of vertebrae in 16 specimens. Vertebral counts followed Hosoya (1983); this included the first four vertebrae with the Weberian apparatus and one fused vertebra of the hypural complex.

### DNA extraction and whole-genome sequencing

To minimize damage to museum specimens, a small clip of pelvic fin tissue (approximately 2 mm × 4 mm) was taken from the right side of 10 *T. lanceolata* individuals for DNA extraction. Fin tissues were lysed overnight at 56 °C in the lysis buffer [10 mM Tris-HCl pH 8.0, 100 mM EDTA, 0.5% SDS containing 0.04 mg/mL of proteinase K solution (Promega)]. DNA was then extracted using the standard phenol–chloroform method and precipitated with isopropanol in the presence of glycogen (final concentration of 0.25 mg/mL; Thermo Scientific) to enhance recovery of short DNA fragments. Purified DNA was quantified with a Qubit fluorometer (Invitrogen), and DNA integrity was assessed using the High Sensitivity DNA Kit on a Bioanalyzer (Agilent).

Long-storage in ethanol is expected to cause DNA degradation through shearing, depurination, and deamination (Raxworthy and Smith 2021). To repair nick and deamination, the extracted hDNA was treated with the NEBNext FFPE DNA Repair v2 Module (NEB) following the manufacturer’s protocol. Repaired DNA was subsequently purified, and fragments < 40-bp were removed using the NucleoSpin Gel and PCR Clean-up (Macherey-Nagel). The resulting DNA was used for whole genome sequencing library preparation using the NEBNext Ultra II DNA Library Prep Kit for Illumina (NEB), performed at half reaction volume with >10 ng DNA input, according to the manufacturer’s instructions. Library quality and concentration were assessed with the High Sensitivity DNA Kit on a Bioanalyzer, and sequenced on an Illumina NovaSeq X Plus (Illumina) or DNB-SEQ (MGI) with 150-bp paired-end reads.

### Bioinformatic analysis

Sequences with short inserts were subjected to trim adapters and low quality bases using fastp v0.20.1 (Chen et al. 2018) with the option ‘-- adapter_sequence=AGATCGGAAGAGCACACGTCTGAACTCCAGTCA -- adapter_sequence_r2=AGATCGGAAGAGCGTCGTGTAGGGAAAGAGTGT -r -q 15 -l 30’. Subsequently, overlapping paired-end reads were merged with FLASH2 software (Magoč and Salzberg 2011) with the option ‘-m 10 -M 65’. Processed reads were mapped to the *T. lanceolata* mitochondrial genome (GenBank accession: KJ589418) (Xu et al. 2016) using BWA-aln v0.7.17 (Li 2013). Small variants were called with the mpileup program implemented in BCFtools v1.13 (Danecek et al. 2021) applying default parameters except for a minimum mapping quality of 20 and a minimum base quality of 15. After converting the variants to VCF format with BCFtools, we selected single nucleotide polymorphisms (SNPs) and filtered out SNPs with QUAL < 20 or depth < 5. Because the analysis focused on the mitochondrial cytochrome *b* (cyt*b*) region, following previous studies (Hashiguchi et al. 2006; Tominaga et al. 2020), the consistency between mapped reads and identified SNPs were manually inspected using the Integrative Genomics Viewer (IGV) (Robinson et al. 2011). For each specimen, mitochondrial haplotype sequences were reconstructed with the consensus program implemented in BCFtools masking positions with coverage depth < 5 as “N”. The phylogenetic position of Vega specimens were compared with the dataset used in the previous studies (Hashiguchi et al. 2006; Tominaga et al. 2020) by constructing a maximum likelihood (ML) tree from partial mitochondrial cyt*b* coding region (1125 bp) using IQ-Tree (Nguyen et al. 2015), with automatic model selection under the vertebrate mitochondrial codon model (-st CODON2) and 1,000 ultrafast bootstrap replicates. Sequences were aligned and curated manually with Aliview software (Larsson 2014), and *T. limbata* was used as the outgroup.

To compare the mappability and characteristics of hDNA with those of high-quality DNA, whole-genome resequencing data from a bitterling fish of the sister genus *Acheilognathus longipinnis* were downloaded (PRJDB15958) (Onuki et al. 2024) and mapped to its mitochondrial genome using the same procedure as described above. This analysis was conducted on a dataset consisting of reads with an original insert size and on a modified dataset in which 60-bp were extracted from one of read pairs to mimic the average insert size of hDNA. This was done by fastp software by specifying the cutting of the 90-bp tail from 150-bp read. Alignment summary statistics were calculated using the idxstats program implemented in SAMtools (Danecek et al. 2021).

To validate the sampling locality, the same procedure was applied to the hDNA of an *A. melanogaster* specimen collected alongside the *T. lanceolata* Vega specimens. The dataset of mitochondrial partial cyt*b* sequence alignments (1125-bp) with a previous phylogeographic study (Nagata et al. 2018) were used to construct a ML tree. As a reference sequence, the *A. melanogaster* mitogenome was newly assembled from a specimen collected in the Kitakami River (Iwate Prefecture). The same procedure was used for DNA extraction and sequencing, but for library preparation, the NEBNext Ultra II FS DNA Library Prep Kit for Illumina (NEB) was used, following the manufacturer’s protocol for a half-volume reaction. *De novo* mitogenome assembly was performed with GetOrganelle v1.8.0.1 (Jin et al. 2020) using short-read sequencing data with the default parameters. The *A. yamatsutae* mitochondrial genome (accession:NC_013712) was used as a seed sequence.

## Results and Discussion

### Morphological identification and preservation conditions of the Vega specimens

Bitterling specimens from the Kanto region, registered as *A. melanogaster*, were first subjected to morphological identification by measuring and counting their diagnostic characteristics (Fig1, Table S1). Of the 44 specimens in lot NRM 8168, 43 were identified as *T. lanceolata* and re-registered under NRM 68053, based on the following characteristics: long barbels (mean 18.8% ± 3.6% of head length: HL), 35–38 lateral line scales complete (mode 36), 9 branched soft rays in dorsal fin, 9–10 branched soft rays in the anal fin (mode 10), a row of black spots along the middle position of the dorsal fin membrane, and absence of colored vertical lines. Only one specimen was identified as *A. melanogaster*, characterized by short barbels (10.7% of HL), lateral line scales complete (35–38), 8 branched soft rays in the dorsal fin, 7 branched soft rays in the anal fin, and traces of long colored vertical line. The presence of *T. lanceolata* in the Kanto region provided an opportunity to investigate its controversial distribution i.e., native or non-native using hDNA analysis.

The application and efficacy of hDNA analysis depend strongly on specimens condition (Raxworthy and Smith 2021). Therefore, we assessed external and internal preservation status non-invasively. Externally, most specimens retained firm bodies with almost no scale loss; only minor damage to the caudal fin was observed in some specimens, and no apparent differences in condition among specimens were noted (Fig. 1, Fig. S1A). Internally, soft X-ray imaging clearly revealed intestines and skeletal structures (Fig. S1B). The presence of guanine in the skin and the absence of bone demineralization indicated that the specimens would have not been exposed to formalin after fixation nor stored long-term in low-concentration ethanol. Collectively, these features demonstrate that the specimens were in exceptionally good condition, providing a reliable basis for genetic analysis of over 100-year-old ethanol-preserved museum collections.

### Degraded but relatively longer hDNA in Vega Collection

To establish an efficient workflow for analyzing hDNA of the well-preserved Vega Collection, we first assessed DNA quality. As expected, the hDNA was degraded with a major fragment size peak around 50-bp and a minor fraction extending beyond 300-bp, yielding 44–705 ng from a dorsal fin clip (Fig. 2A, B). Although fragmentation was pronounced, yield and fragment length were higher than those typically recovered from historical formalin-fixed specimens, which often show low yields and more severe degradation (Hahn et al. 2022).

**Figure 2.**
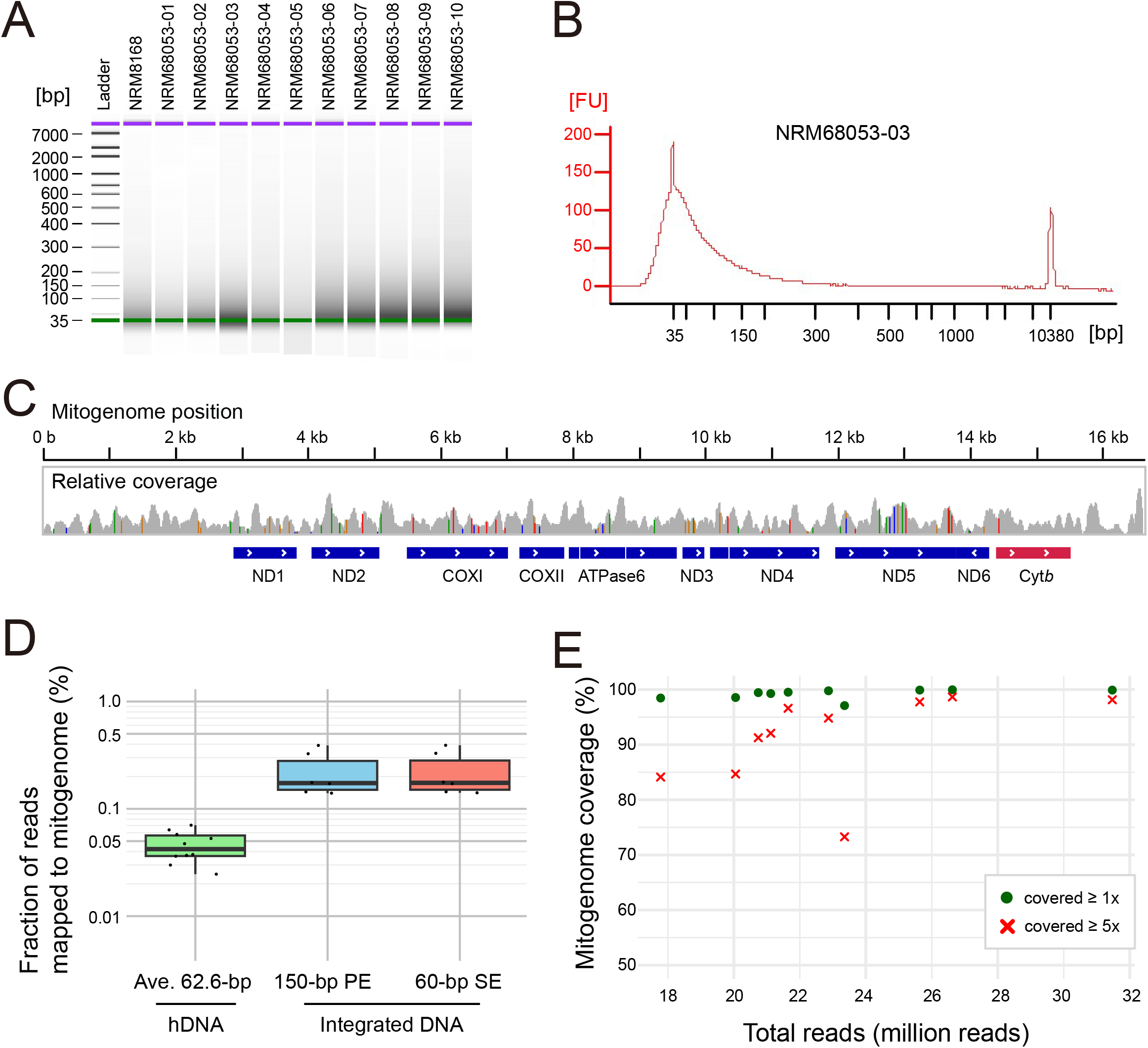
Characteristics of hDNA from 140-year-old ethanol-preserved specimens. (A) Electropherogram showing the fragment length distribution of DNA extracted from Vega specimens. (B) Representative Bioanalyzer electropherogram. (C) Representative distribution of mapped reads to the *T. lanceolata* mitogenome and inferred SNPs, visualized in the Integrative Genomics Viewer. Gray bars indicate the relative coverage per site (maximum depth 98x). SNPs are indicated by colored bars corresponding to the alternative base: A (green), T (red), G (brown), or C (blue). Blue and red boxes indicate mitochondrial coding genes, with arrows denoting gene orientation. (D) Log-scaled fraction of reads mapped to the conspecific mitogenome reference, comparing hDNA (*T. lanceolata* from the Vega Collection) and with integrated DNA (freshly-ethanol-fixed fin tissue from, *A. longispinnis*). (E) Ratio of mitogenome-covered regions relative to total reads for each Vega specimen. Green dots and red crosses represent regions covered ≥ 1x and ≥ 5x, respectively.

The relatively high yields and broader fragment size distribution prompted us to repair DNA damage before and subsequently remove short molecules (∼40 bp) using column-based purification, thus improving sequencing efficiency and reducing unwanted C/T substitutions caused by deamination.

To evaluate the efficacy of sequencing hDNA, we compared the mappability of short reads to the *T. lanceolata* mitochondrial genome with that of high-quality DNA extracted from the 99%-ethanol fixed fin tissue of *A. longipinnis*. For hDNA from the Vega specimens, 0.02–0.07% of reads mapped to the conspecific mitogenome reference, approximately fivefold lower than the mapping rate of integrated DNA, even after equalizing read length (Fig. 2D). Nevertheless, analysis of over 10 million reads provided coverage of above 97% of the mitogenome with at least one read, and nearly complete coverage at ≥ 5-fold depth was achieved with ∼30 million reads (Fig. 2E). These results indicate the sequencing depth required and demonstrate that the 140-year-old Vega specimens are suitable for genomic analysis.

### Unexpected genetic similarity between modern and the Vega specimens

To test the translocation hypothesis from the Kinki to the Kanto regions, we examined the phylogenetic position of the Vega specimens. A maximum likelihood (ML) tree was constructed using partial mitochondrial cytochrome *b* (cyt*b*) sequences, as in a previous phylogeographic study (Tominaga et al. 2020). All 10 Vega specimens analyzed shared an identical haplotype, belonging to a clade (Clade A) widely distributed across the Kinki, Sanyo, Shikoku, and Kanto regions (Fig. 3). Similar mitochondrial haplotypes were also observed in modern specimens from multiple locations of the Kanto region and several sites on the western side of the Central Highland, with no apparent sequence divergence (Fig. 3).

**Figure 3.**
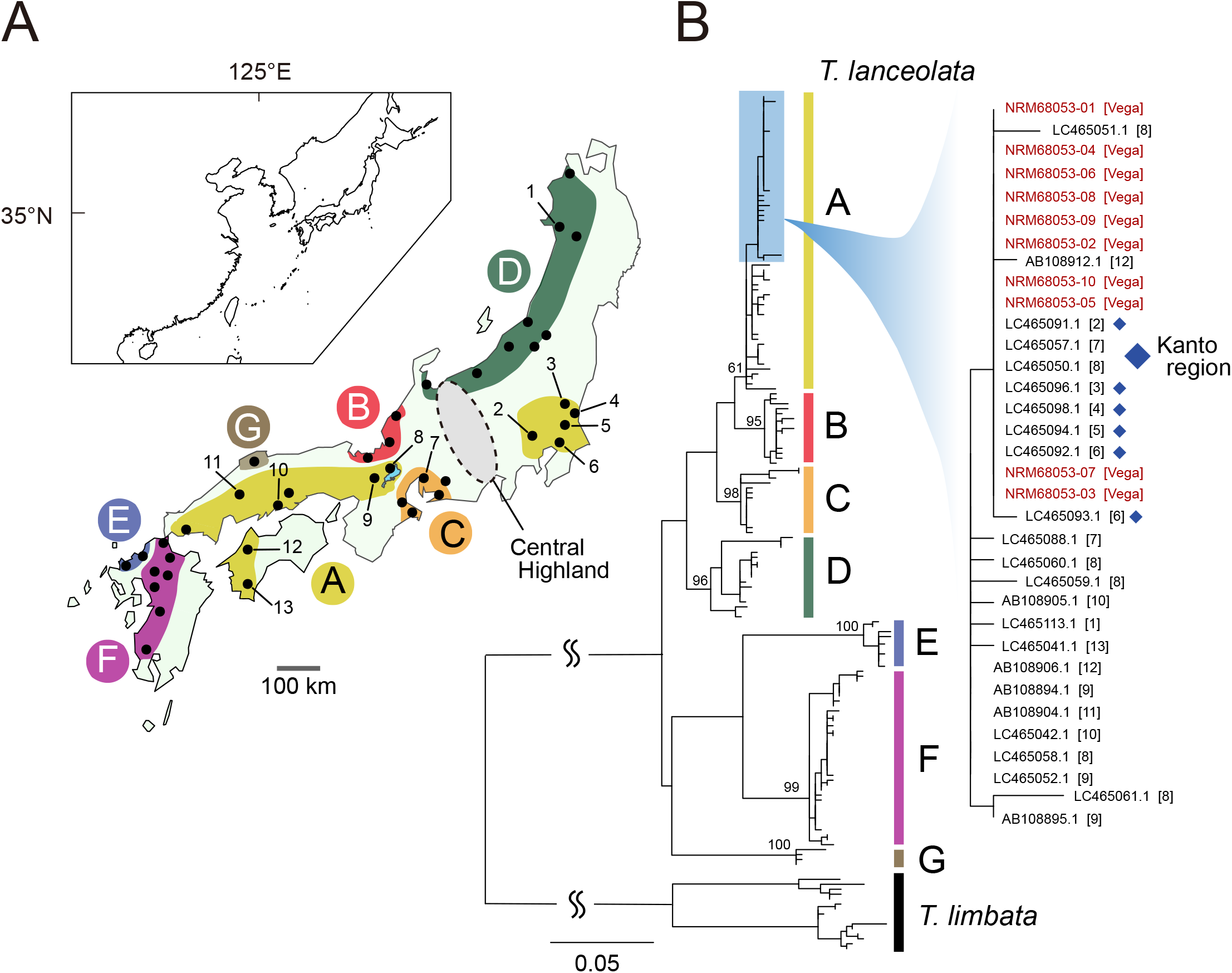
Mitochondrial genetic similarity between Vega and modern specimens from the Kanto region. (A) Map showing the locations of analyzed specimens and the approximate distribution of clades A–E (adapted from Tominaga et al. 2020). Black points indicate sampling sites, with numbered localities (1–13) representing detection of mitochondrial haplotypes similar to those of the Vega specimens. (B) ML tree of *T. lanceolata* based on partial mitochondrial cyt*b* sequences. A subclade containing the Vega specimens is magnified, with sample IDs shown alongside locality codes. Vega specimens are highlighted in red, and modern specimens from the Kanto region are marked with blue diamonds. Numbers on major nodes indicate bootstrap values.

When analyzing historical specimens, the possibility of erroneous locality records must be considered (e.g., Cong et al. 2021). In this study, the locality of the analyzed Vega specimens was validated by the presence of a co-collected bitterling identified morphologically as *A. melanogaster*, a species endemic to the Pacific coast side of eastern Japan. Genetic analysis of mitochondrial sequences further confirmed this identification, showing a haplotype consistent with a group distributed around the Kanto region (Locality I; Fig. S2). These results indicate that the mitochondrial genetic similarity of *T. lanceolata* across the Central Highland was already established by the 1870s.

The genetic similarity of *T. lanceolata* across the Central Highland could reflect past large-scale domestic translocations from Kinki to Kanto. Intended for fishery enhancement, such translocations have been documented since at least 1918 and have continued since then, occasionally causing the unintended translocation and establishment of several non-subjective fish species such as *Ischikauia steenackeri, Opsariichthys uncirostris* and *Rhynchocypris lagowskii steindachneri* (Yanai et al. 2008; Nishida et al. 2025). Notably, these translocations have occurred from Lake Biwa to Lake Kasumigaura, a large lake connected to the Tone River, which flows across a large area of the Kanto Plain (Yanai et al. 2008). However, Vega specimens analyzed here were collected before these large-scale fishery efforts, suggesting that the widespread distribution of mitochondrial haplotypes across the Central Highland may instead reflect unanticipated natural dispersal, as observed in Japanese medaka (*Oryzias latipes*) (Katsumura et al. 2018). Alternatively, it may result from undocumented historical translocations before 1879. Future analyses using comprehensive genome data from both historical Vega specimens and modern individuals across multiple localities will help to resolve these scenarios, clarify long-term changes in genetic diversity, and illuminate the long-term evolutionary dynamics following potential translocations.

## Conclusion

In this study, we assessed the hDNA quality in relation to specimen preservation and established a workflow for genetic analysis of ethanol-preserved specimens collected during the Vega Expedition 140 years ago. Our results highlight the value of the Vega Collection as its well-preserved condition and demonstrate the potential application of conservation museomics to resolve complex population histories, as exemplified by *T. lanceolata*. Populations suspected to be non-native or impacted by genetic pollution are often deprioritized in conservation strategies; therefore, accurate reconstruction of population histories is critical for effective management. The workflow developed here is broadly applicable not only to other Vega specimens but also to historical ethanol-preserved collections, thereby enhancing the scientific value of archival specimens.

## Data availability

The sequence data generated and analyzed in this study were deposited in the Short Read Archive (SRA) under accession number of PRJNA1338267.

## Acknowledgement

We thank Bo Delling (NRM) for specimen loans and information on storage conditions. We thank the Center for Advanced Technical and Educational Supports, Faculty of Agriculture, Kyushu University for the use of facilities. We thank Katsutoshi Watanabe (Kyoto University) for discussion on data interpretation. This work was supported by JSPS KAKENHI Grant Number 15H05234 to YT, and 23K05899 to SK, and 25K02327 to TM.

## Author contribution

Surveys for the Vega Collection were performed by SK, YT, TF, and KH; Morphological analysis were performed by SK; hDNA experiment and data analysis were performed by KT and TM; Interpretation of data/results by KT, SK, JK and TM. Writing of the first draft of the manuscript by SK and TM. Funding acquisition by YT, SK, and TM; Project administration by SK, JK, and TM; All authors revised the manuscript and accepted the final version for publication.

## Figure Legends

**Figure S1.**
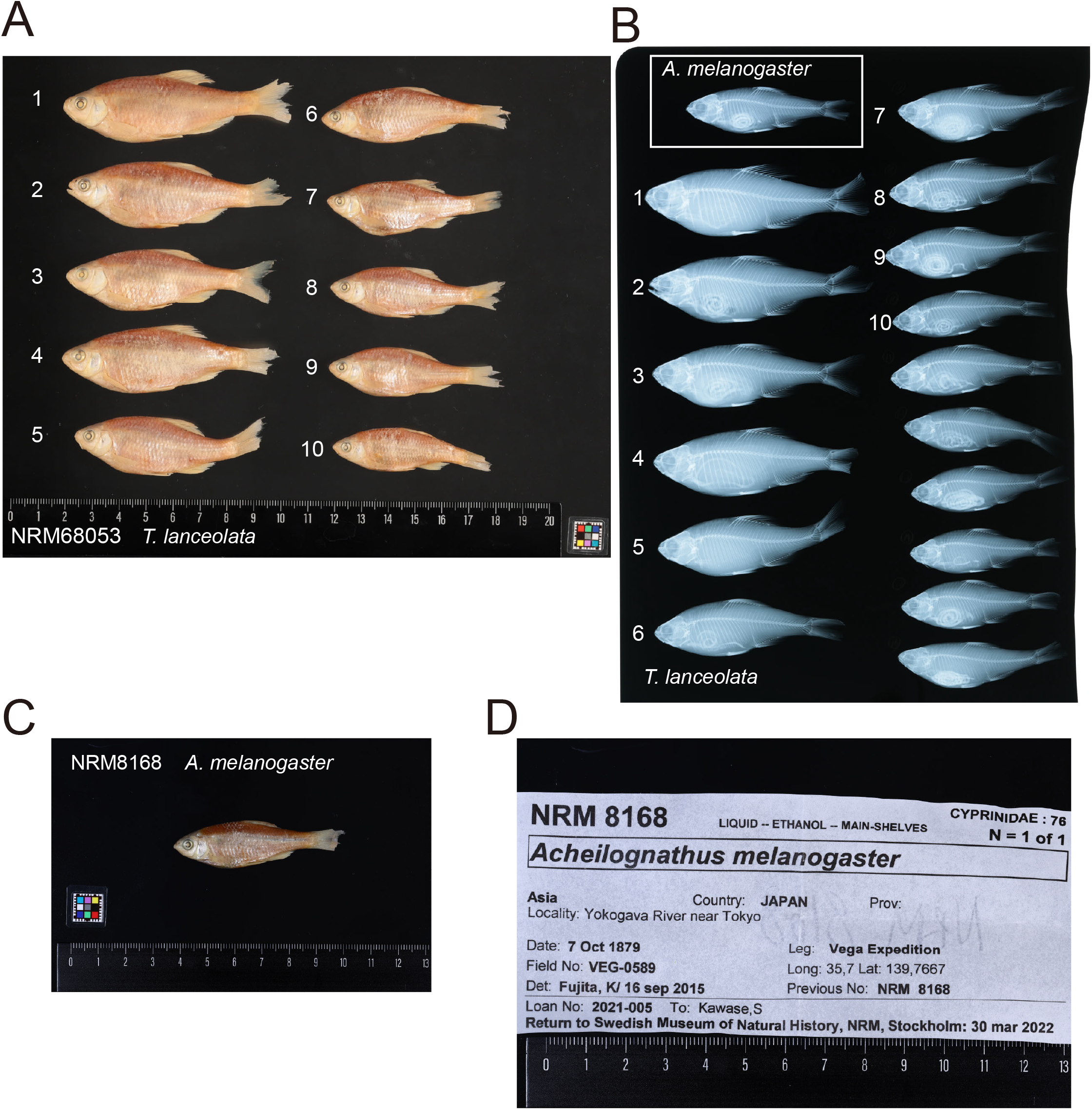
Preservation status of analyzed bitterling specimens in the Vega Collection. (A) Lateral view of the ten sequenced *T. lanceolata* specimens, numbered according to soft X-ray images. (B) Soft X-ray image of *T. lanceolata* and *A. melanogaster* specimens, showing well-preserved bones, rays, and digestive tracts. (C) Lateral view of *A. melanogaster*. (D) Revised specimen label with detailed original collection records.

**Figure S2.**
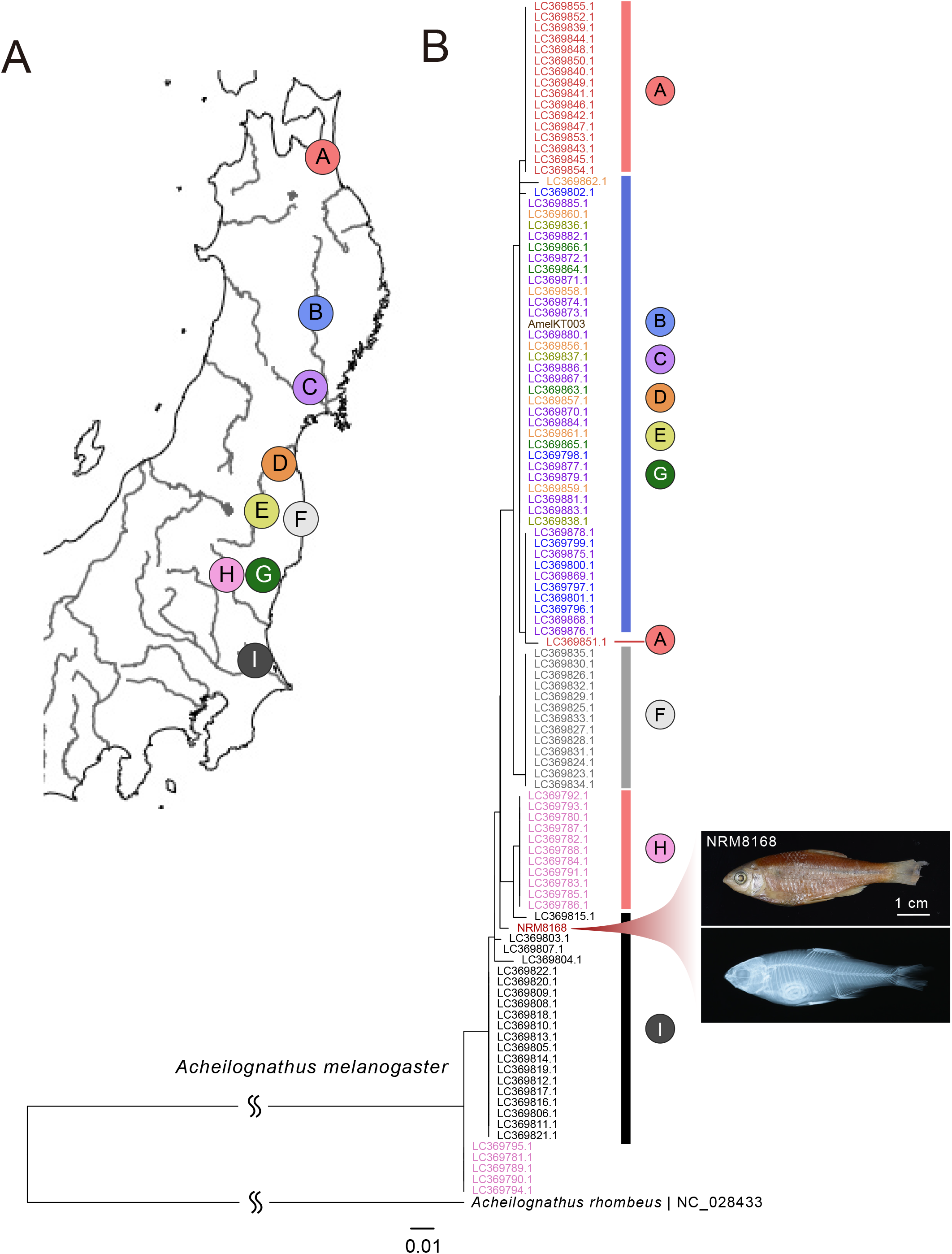
Validation of the sample locality using a co-sampled bitterling fish, *A. melanogaster*. (A) Sampling localities of the re-analyzed specimens used in Nagata et al. 2018. (B) ML tree based on mitochondrial cyt*b* sequences, showing that the *A. melanogaster* specimen from the Vega Collection, recorded as locality of “Yokogawa River near Tokyo” clusters with specimens from the Kanto plain (Locality I). Accession IDs derived from Nagata et al. 2018 are color-coded to sampling locality.

**Table S1. Summary of morphological characteristics of 30 *T. lanceolata* and 1 *A. melanogaster* specimens from the Vega Collection**. Scale counts, based on soft X-ray images, were performed on 16 *T. lanceolata* and 1 *A. melanogaster* (denoted with *). SL: standard length; HL: head length; SD: standard deviation.

